# Impaired microglia-mediated synaptic pruning in the nucleus accumbens during adolescence results in persistent dysregulation of familiar, but not novel social interactions in sex-specific ways

**DOI:** 10.1101/2023.05.02.539115

**Authors:** Julia M. Kirkland, Erin L. Edgar, Ishan Patel, Ashley M. Kopec

**Author notes:** Corresponding Author: J. M. Kirkland, 43 New Scotland Ave., Albany, NY 12208.

## Abstract

Evolutionarily conserved, peer-directed social behaviors are essential to participate in many aspects of human society. These behaviors directly impact psychological, physiological, and behavioral maturation. Adolescence is an evolutionarily conserved period during which reward-related behaviors, including social behaviors, develop via developmental plasticity in the mesolimbic dopaminergic ‘reward’ circuitry of the brain. The nucleus accumbens (NAc) is an intermediate reward relay center that develops during adolescence and mediates both social behaviors and dopaminergic signaling. In several developing brain regions, synaptic pruning mediated by microglia, the resident immune cells of the brain, is important for normal behavioral development. In rats, we previously demonstrated that microglial synaptic pruning also mediates NAc and social development during sex-specific adolescent periods and via sex-specific synaptic pruning targets. In this report, we demonstrate that interrupting microglial pruning in NAc during adolescence persistently dysregulates social behavior towards a familiar, but not novel social partner in both sexes, via sex-specific behavioral expression. This leads us to infer that *naturally occurring* NAc pruning serves to reduce social behaviors primarily directed toward a familiar conspecific in both sexes, but in sex-specific ways.

## INTRODUCTION

Social interactions are a crucial part of existence across the lifespan (Haslam et al., 2021; Schweinfurth, 2020; Siracusa et al., 2022; Yang et al., 2016) in most species. Studying the development of normative social behavior is critical for recognizing the evolutionary goals of sociability (Cavigelli & Caruso, 2015; Li et al., 2021) and the vulnerability conferred by abnormal social behavior (Almquist et al., 2018; Burke et al., 2017; Cameron et al., 2017; Hodges et al., 2017). Moreover, a better understanding of the neural mechanisms of social development may be therapeutically relevant, as positive social support is shown to improve outcomes in cognitive aging (Crimmins, 2020; Frey et al., 2021; Leblanc & Ramirez, 2020)], recovery post-cardiac events (McNeal et al., 2014) and recovery from substance abuse (Boisvert et al., 2008; Venniro et al., 2018), among others. Conversely, negative social experiences are predictive of negative outcomes both mentally (Al Omran et al., 2022; Von Frijtag et al., 2002; Weintraub et al., 2010) and physically (Kang et al., 1998; Okuda et al., 2022; Schneider et al., 2016). Adolescence is an evolutionarily conserved period during which reward-related behaviors, including social behaviors, develop via developmental plasticity in the mesolimbic dopaminergic ‘reward’ circuitry of the brain (Brenhouse & Schwarz, 2016; Manduca et al., 2016). The nucleus accumbens (NAc) is an intermediate reward relay center that develops during adolescence (Bayassi-Jakowicka et al., 2021; Huppe-Gourgues & O’Donnell, 2012; Huppé-Gourgues & O’Donnell, 2012; Kopec et al., 2018; Martz et al., 2022; Orihuel et al., 2021) and mediates both social behaviors and dopaminergic signaling (Halbout et al., 2019; Manduca et al., 2016). In several developing brain regions, complement system proteins (e.g., C3) facilitate synaptic pruning by binding to complement receptor 3 (CR3) on microglia, the resident immune cells of the brain (Grabert et al., 2016; Schafer et al., 2012; Soteros & Sia, 2022). In rats, we demonstrated that C3-microglial synaptic pruning also mediates NAc and social development during sex-specific adolescent periods pre- to early adolescence in females (postnatal day (P)22-30) and early to mid-adolescence in males (P30-P40), and via sex-specific synaptic pruning targets (dopamine D1 receptors in males, but not females) (Kopec et al., 2018). Despite divergent timing and targets, inhibiting microglial pruning in the NAc increased adolescent social play, a developmentally regulated social behavior, in both sexes. Whether inhibiting NAc pruning during sex-specific pruning periods would increase sociability persistently into adulthood, and if its behavioral manifestation would remain the same in both sexes, is unknown. In the following experiments, we seek to identify how sex-divergent neuroimmune activity in the NAc impacts normal social development. We hypothesized that inhibiting sex-specific NAc pruning during adolescence would increase sociability in sex-specific ways in adulthood. To test this hypothesis, we inhibited C3-microglial pruning in the NAc during each sex’s respective pruning period with a single, bilateral microinjection of neutrophil inhibitory factor (NIF), a highly specific competitive CR3 antagonist (Muchowski et al., 1994; Smirnov et al., 2023). Rats were permitted to age into adulthood, then assessed through a social battery. We found that interfering with natural NAc pruning at sex-specific timepoints increased pro-social behavior primarily toward familiar, but not novel, social partners, in sex-specific ways.

## METHODS

### ANIMALS

Adult male and female Sprague-Dawley rats were purchased for breeding (Envigo, Indianapolis, Indiana) and group-housed with ad-libitum access to food and water in cages with cellulose bedding changed twice weekly. Colonies were maintained in a 12:12 light: dark cycle (lights on at 07:00) in a temperature and humidity-controlled room. Litters were culled to a maximum of 12 pups per dam between postnatal day 2 (P2) and P5. At P21 pups were weaned into same-sex pairs. Rats were housed in the Animal Resources Facility in Albany Medical College and all animal work was approved by the Institutional Animal Care and Use Committee at Albany Medical College.

### BILATERAL STEREOTACTIC INFUSION INTO THE NAC

Stereotactic procedures were performed experimentally determined in Kopec et al. 2018. Male (P30) and female (P22) rats were maintained under isoflurane anesthesia for the entire surgical procedure (2–3%; KENT SCIENTIFIC; Livermore, CA). The scalp was cut midsagittally and Bregma was marked, after which two bilateral holes drilled at *AP + 2*.*25 mm, ML ± 2*.*5 mm, DV −5*.*75 mm* coordinates in P30 males, *and AP + 2*.*7 mm, ML ± 2*.*4 mm, DV -5*.*55 mm* in P22 females. A Hamilton syringe (Hamilton #7105; Reno, NV) was lowered to depth at a 10° angle and left in place for 1 min. NIF (1x reconstitution, 200 μg/mL NIF; R&D Systems; Minneapolis, MN) or vehicle (sterile PBS) was injected at a rate of 50 nL/min (60 ng in 300nL in P30 males, 50ng in 250nL in P22 females). The syringe was left in place for 5 min post-infusion, and then retracted and the procedure repeated on the other hemisphere. The syringe was thoroughly cleaned with cleaning solution (Hamilton cleaning concentrate, Hamilton; Reno, NV) and distilled water before being used for the next surgery. Saline was used to wet the scalp, and then the wound was closed with surgical staples and coated topically with Bupivacaine. An injection of Ketophen (5 mg/kg) was administered subcutaneously and the animal was placed in a clean cage with a food pellet and gelatinized water for recovery. Animals were re-paired prior to being returned to the animal facility. Animals received Ketophen (5 mg/kg) for two additional days following surgery (3 total injections, once per day), and staples removed 10 days post-surgery. Rats were left undisturbed until behavioral testing in adulthood.

### BEHAVIORAL TESTS

Experimental rats (∼P90) and their sex- and age-matched novel conspecifics were acclimated to experimenter handling and the behavior room for ∼5 minutes per animal for 3 weekdays preceding behavior tests. The order of behavioral tests was counterbalanced and one behavior test per day was performed. All behavior tasks were recorded using ANY-Maze software and the first 5 minutes were hand-coded by blinded experimenters for analysis in Solomon Coder software (Andras Peter; solomoncoder.com). ***<u>Open field test (OF):</u>*** experimental rats were placed in an empty open field box for 10 minutes with overhead recording. The time spent in the center (inner zone) was quantified. ***<u>Social choice tasks:</u>*** (*social novelty preference (SNP) and social versus object choice (SVO)*) experimental rats were first acclimated (10mins) in the empty three-chamber box, a black plexiglass behavior box consisting of three compartments: an experimental compartment flanked by two smaller compartments where stimuli (social and nonsocial) are securely housed. Stimuli are separated from the experimental compartment by a transparent plastic wall with several small holes not large enough for an animal to move across. Choice tasks were scored for time spent in either exploratory zone (i.e., next to either stimulus). The percentage of exploration was calculated using the time spent exploring the stimulus as a function of the total time spent in the corresponding half of the experimental compartment (e.g., the time spent exploring the nonsocial object divided by the total time spent on the nonsocial object half of the apparatus; **Supp. Fig. 1A)**. ***<u>Novel and familiar social interaction tests (novel and familiar social interaction)</u>*** were performed in clean, empty rat housing cages. Rats were acclimated for ∼10mins to the interaction cages prior to each experiment. Novel social interaction was performed by placing an experimental animal and a novel age- and sex-matched conspecific in the same cage for 10 minutes. Familiar social interaction was performed by placing two familiar cage mates that had undergone the same adolescent manipulation in the same cage for 10 mins. Cage mates were separated for ∼20min prior to test. Interaction behaviors were quantified by blinded hand-coding coding of time spent in four social behavior phenotypes. 1. Active social behavior is defined as the rat facing, and physically in contact with, the conspecific. 2. Passive social behavior occurs when the rat is facing the conspecific with no physical contact. 3. Nonsocial contact is when the rat is engaged in physical contact with the conspecific but is not facing the animal. 4. Nonsocial attention occurs if the rat is not facing nor in physical contact with the conspecific **(Supp. Fig. 1B)**.

### STATISTICS

For all behavior tasks, Student’s two-tailed t-test (p ≤0.05) was used to compare NIF-treated to Vehicle-treated animals of the same sex. Because surgeries were performed at different ages for each sex, corresponding to their respective NAc pruning periods, we could not compare directly between the sexes. Outliers were identified using the ROUT method (Q=1%) and eliminated from analysis. For cages of three, only one pairing of data was included for the familiar interaction task, as post-hoc analysis indicated significantly different responses to the familiar rat when it was used for a second test. Correlations for open field results were calculated using Pearson’s r. Statistically significant r values were compared between NIF-treated and PBS-treated animals via Fisher’s r-to-z transform to identify any significant change resulting from inhibition of pruning in the NAc, and a Benjamini-Hochberg correction was used to account for multiple comparisons. A p-value of less than 0.05 is defined as statistically significant. Statistics were performed using GraphPad Prism Version 9.5.0 and vassarstats.com.

## RESULTS

### Inhibiting adolescent NAc pruning does not change social vs. object choice behavior in either sex

Social vs. object choice tests are a widely used assay for quickly and simply assessing sociability (Wöhr & Scattoni, 2013). Rodents have a natural preference for novel social stimuli over novel object stimuli, and under normal conditions will thus spend more time exploring the novel social stimulus. This was indeed the case in both males and females of both experimental groups, but contrary to our hypothesis, inhibiting pruning during adolescence did not impact the behavioral parameters we measured during social vs. object (**Fig 2**) in either sex.

**Fig 1.**
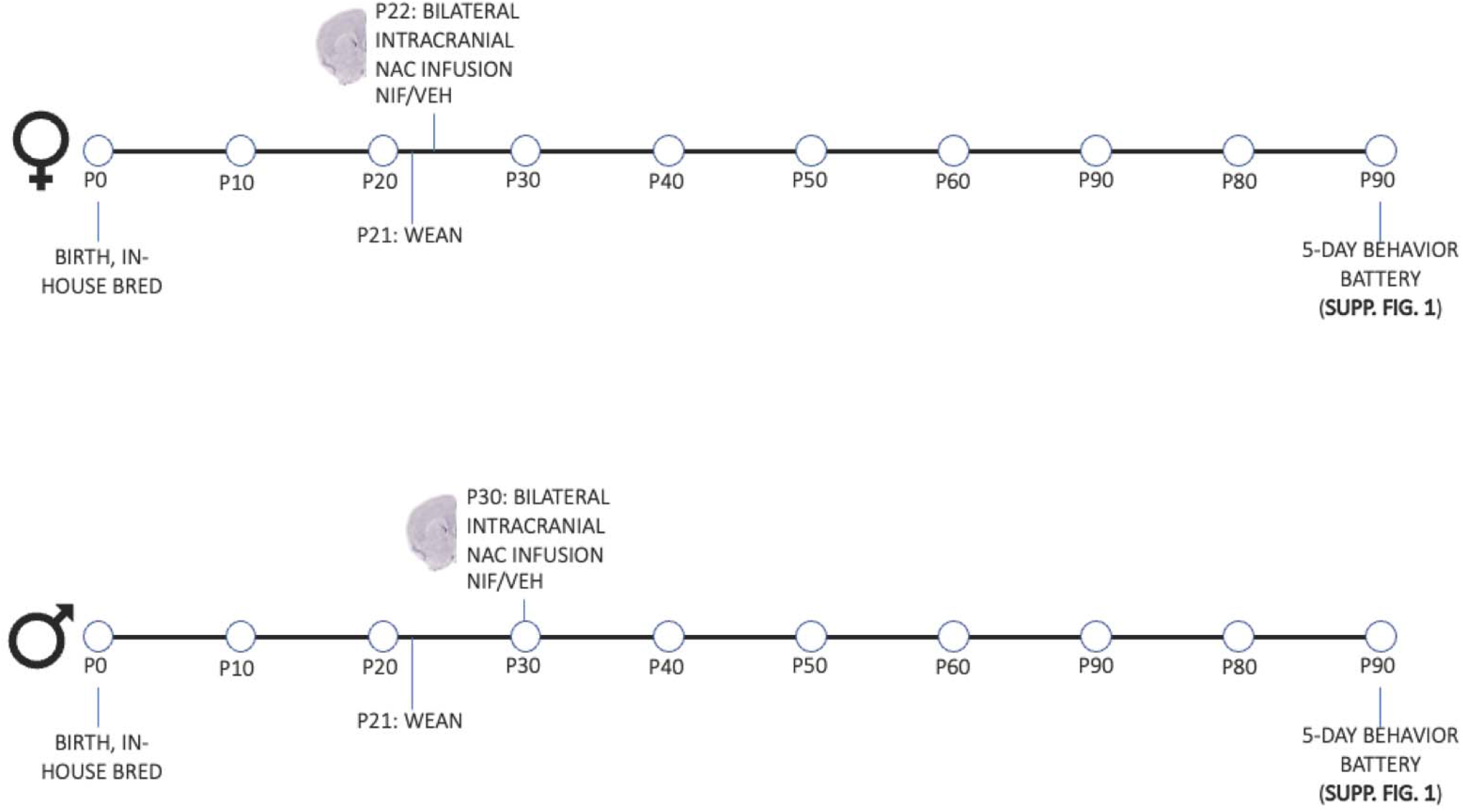
Experimental Design.

**Fig 2.**
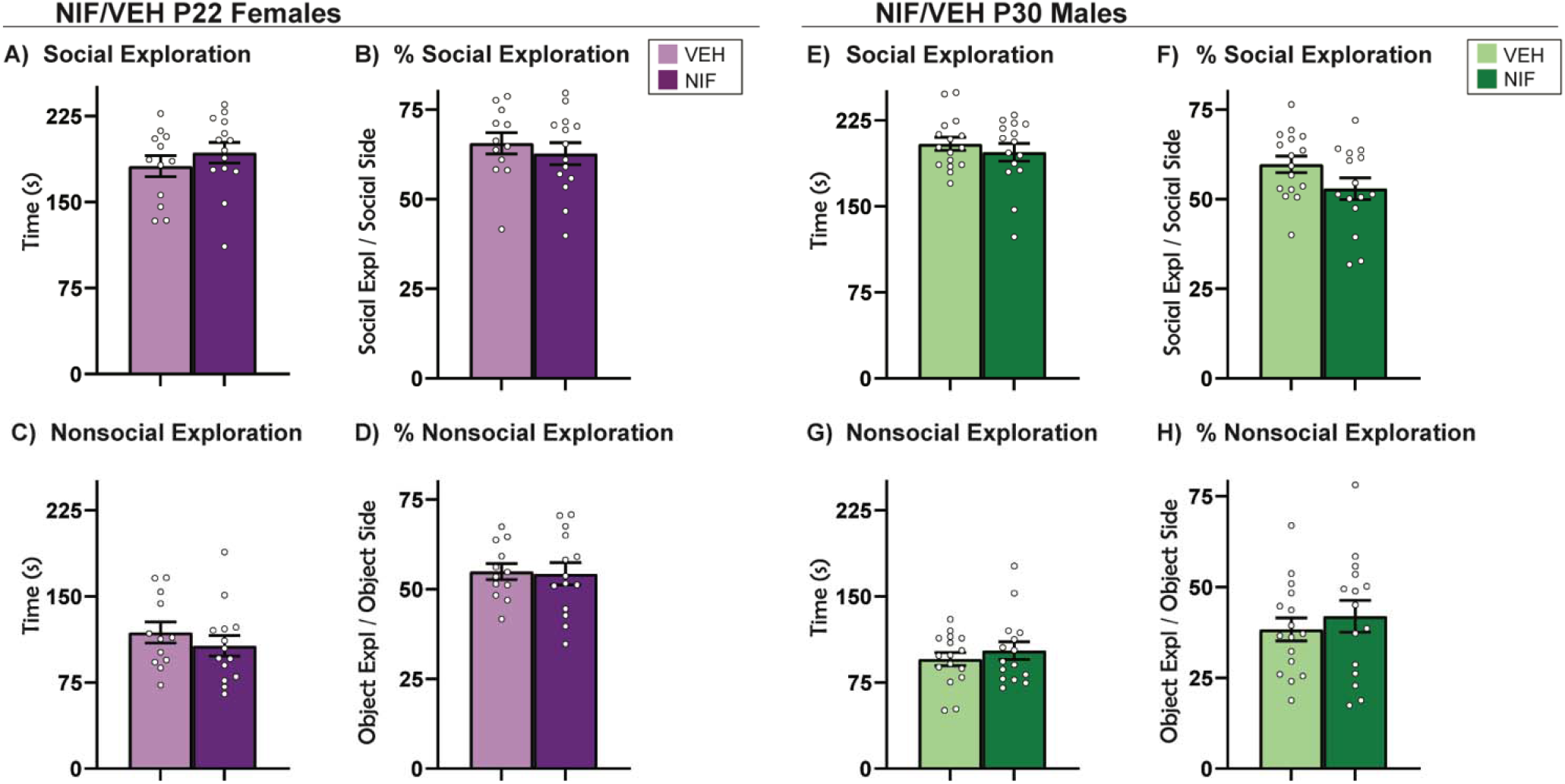
Social vs Object Choice. Inhibiting pruning in the NAc during adolescence did not impact the behavioral parameters measured in the Social vs. Object Choice tests in (A-D) females or (E-H) males. Histograms represent average ± standard error of the mean. Student’s two-tailed t-test. If ^*^ then p< 0.05; n=12-17/sex/group

### Inhibiting adolescent NAc pruning alters behavior towards a familiar partner during social novelty preference task in males, but not females

A relatively new variant of social choice tests is Social Novelty Preference (SNP) (Smith et al., 2015; Smith et al., 2018), which provides the experimental animal with the opportunity to explore familiar and novel social stimuli in opposing sides of the behavior chamber. Novel and familiar social stimuli serve different purposes in a social network and evoke different social behaviors (Beery & Shambaugh, 2021). Rodents have a natural preference for social novelty over social familiarity, and under normal conditions will thus spend more time exploring the novel social stimulus. In females, inhibiting NAc pruning during adolescence did not change behavior in this test (**Fig. 3A-D**). In males, inhibiting NAc pruning during adolescence did not change the total amount of time spent exploring either stimulus (**Fig. 3E, 3G**), but did increase the time spent exploring the familiar stimulus as a percentage of time in the familiar half of the arena (%exploration) **(Fig 3F)** (t(30)=2.353, p=0.0253).

**Fig 3.**
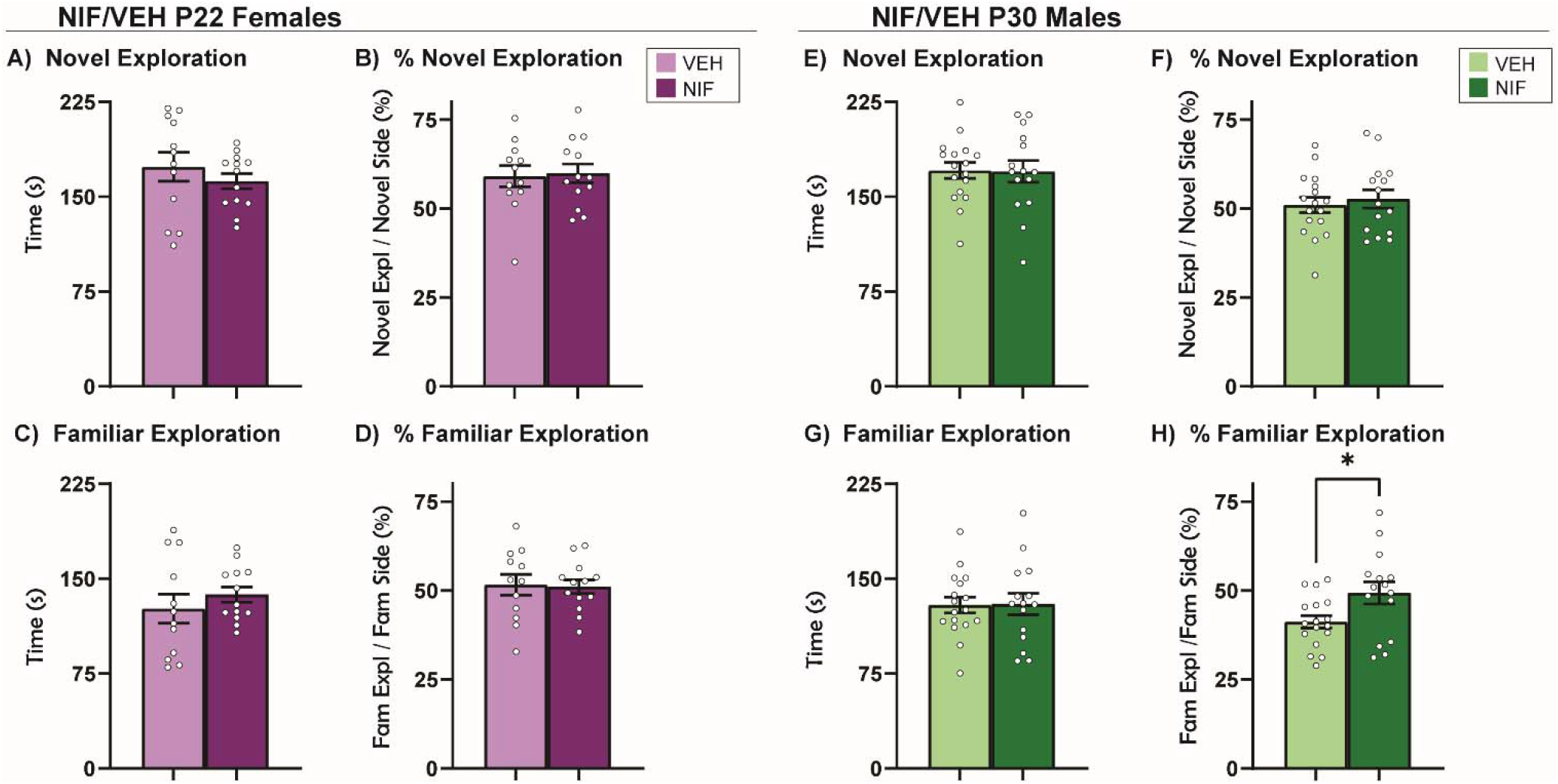
Social Novelty Preference. (A-D) In females, inhibition of NAc pruning during adolescence did not significantly regulate the behaviors measured in the Social Novelty Preference test. (E, F) In males, inhibition of NAc pruning during adolescence did not significantly influence exploration time or %exploration of a novel partner or (G) change time spent exploring familiar partner in males. (H) Inhibiting NAc pruning during adolescence in males significantly increased %exploration of a familiar partner. Histograms represent average ± standard error of the mean. Student’s two-tailed t-test. If ^*^ then p< 0.05; n=12-17/sex/group.

### Inhibiting adolescent NAc pruning increases passive social interaction with a novel animal in males, but not females

To this point, the data has not elucidated any changes in female social choice after NAc pruning. In males, the data shows subtle shifts in familiar-directed exploratory behavior in males, but no changes in behavior towards a novel social conspecific. Novel social interaction is a more naturalistic assessment of direct social interaction in rats (Kraeuter et al., 2019a; Schiavi et al., 2022). We hypothesized that a long-term consequence of inhibiting NAc pruning would be a persistent increase in active social behavior towards a novel conspecific. In males and females, four phenotypes of social behavior were quantified: active social interaction, passive social interaction, nonsocial contact, and nonsocial attention (see **Supp. Fig. 1** and Methods for more detail). There was no significant change in behavior towards a novel partner in females as a result of inhibiting NAc pruning during adolescence (**Fig 4A-D**). Inhibiting NAc pruning in males during adolescence caused the rats to engage in more passive social interaction than control counterparts (t(29)=2.218, p=0.0346) (**Fig 4F)**, but no other changes were detected in active social interaction, nonsocial contact, or nonsocial attention (**Fig. 4E,G,H**).

**Fig 4.**
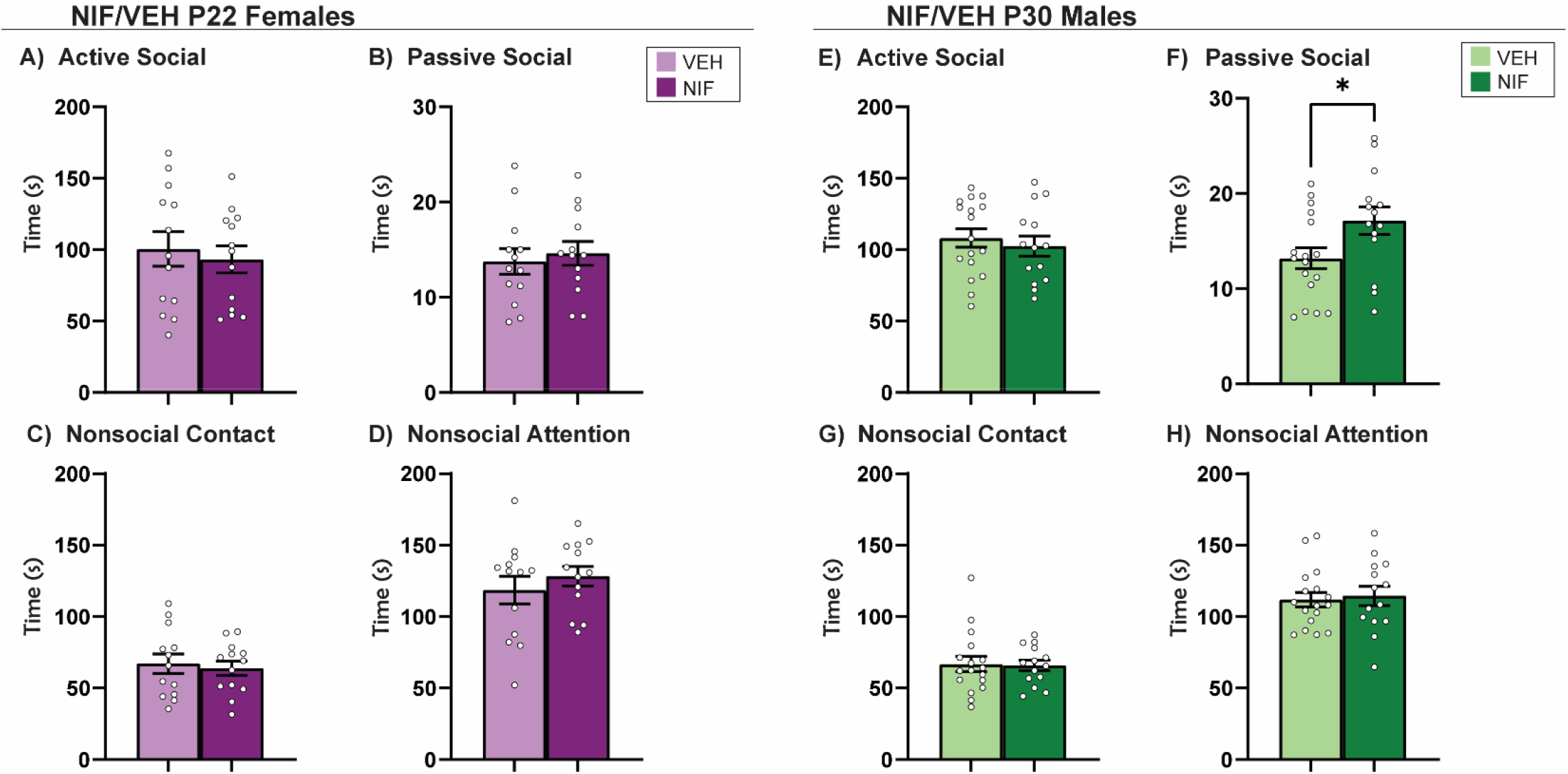
Novel Social Interaction. Inhibiting pruning in the NAc during adolescence did not impact novel social interaction in (A-D) females. In males, inhibiting pruning in the NAc during adolescence does not impact active social interaction (E), nor nonsocial interactions (G-H) with a novel partner. However, time spent engaged in passive social interaction (F) is significantly increased after inhibiting pruning in the NAc. Histograms represent average ± standard error of the mean. Student’s two-tailed t-test. If ^*^ then p< 0.05; n=13-17.

### Inhibiting adolescent NAc pruning increases prosocial behaviors with a familiar cage mate in both males and females, in sex-specific ways

The data thus far have not supported our hypothesis that there will be increased sociability in NIF-treated females, and in males the effects of NIF treatment have been subtle. To fully assess the variety of social behaviors that adult rats display, rats were paired with their cage mates for familiar social interaction, a less commonly used assessment of direct social interaction in cohabitating rats (Varlinskaya & Spear, 2008). In our studies, cage mates have undergone the same adolescent manipulation (e.g. both vehicle- or both NIF-treated), and both animals in the test are scored for the same phenotypes measured in Novel social interaction. Inhibiting adolescent NAc pruning in females increased active social behavior (*t*(20)=2.426, p=0.0248) and reduced nonsocial contact (*t*(20)=2.559, *p*=0.0187) with a familiar cage mate in adulthood (**Fig 5A, Fig 5C)**. Inhibiting NAc pruning during adolescence in males, on the other hand, increased nonsocial contact (*t*(28), p=0.0016) and reduced nonsocial attention (*t*(28)=2.247, *p*=0.0327) (**Fig 5G, Fig 5H**).

**Fig 5.**
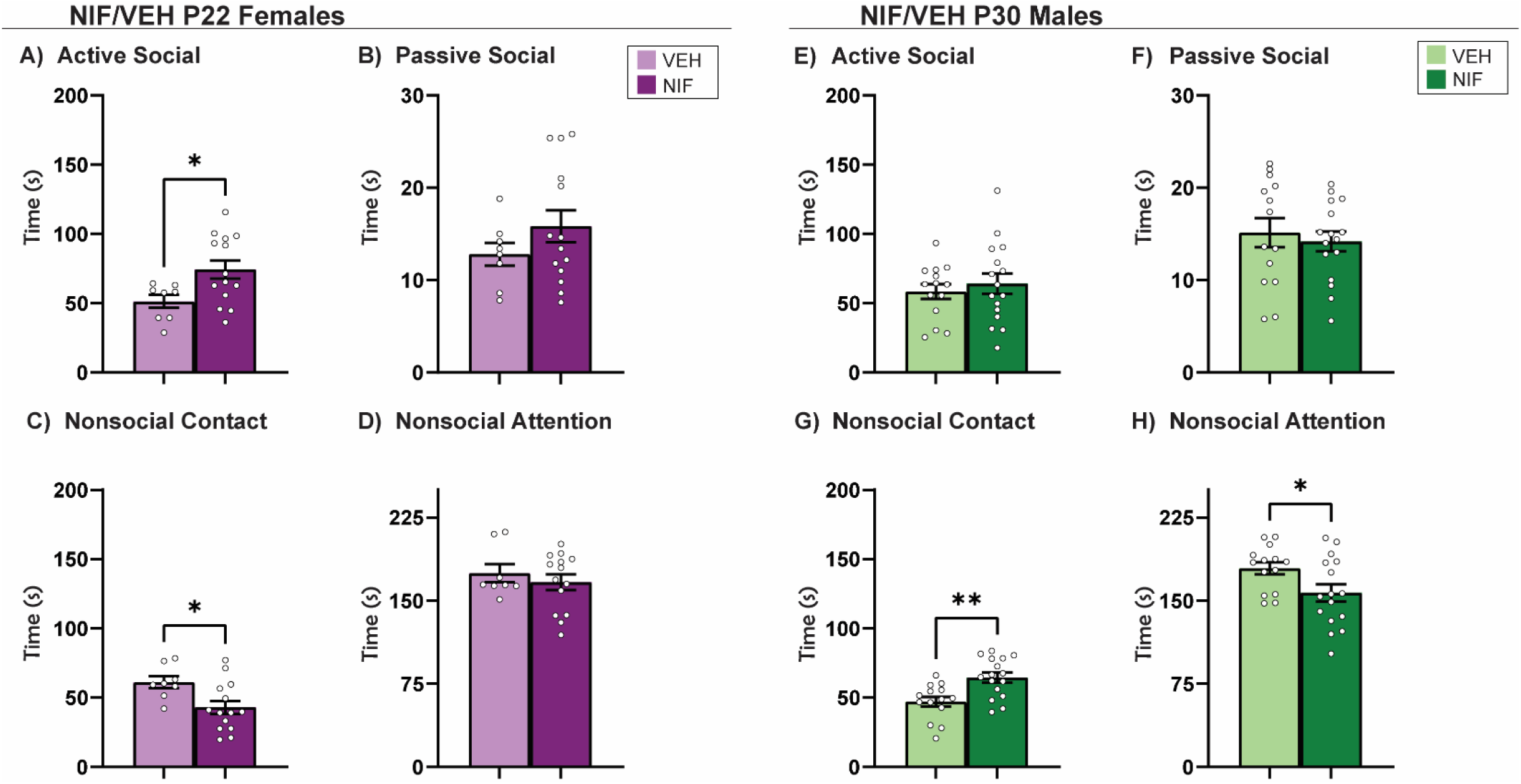
Familiar Social Interaction. (A) Inhibiting NAc pruning during adolescence significantly increased active social attention and (C) significantly decreased nonsocial contact in females. (B, D) No significant changes were seen in passive social or nonsocial attention in females. (E-F) Inhibiting NAc pruning during adolescence did not significantly change active or passive social attention in males. (G) However, inhibiting NAc pruning during adolescence in males significantly increased nonsocial contact and (H) significantly decreased nonsocial attention. Histograms represent average ± standard error of the mean. Student’s t-test. If ^*^ then p< 0.05, if^**^ p< 0.01; n=8-16.

### Inhibiting adolescent NAc pruning decreases anxiety-like/increases exploratory behaviors in males, but not females, in the open field

Open field is a common task designed to assess anxiety-like and/or exploratory behaviors in rodents (Falco et al., 2014; Kraeuter et al., 2019b). We utilized this task to assess whether altered social behavior may be driven by changes in generalized anxiety-like and/or exploratory behaviors in rats that have had NAc pruning inhibited during adolescence. NIF-treated males spent significantly increased time in the inner zone of the open field compared to control males (t(31)=2.552, *p*=0.0159) **(Fig 6B)**, suggesting they were expressing less anxiety-like behaviors and/or increased exploratory behaviors. There were no significant differences between groups in females **(Fig 6A)**. To begin to determine whether a change in anxiety-like/exploratory behavior would be a critical factor in the social differences we observed, we calculated the correlations between open field behavior with the other social metrics presented herein. In females, there were two significant correlations between open field behavior and social behaviors in females (NIF Novel nonsocial attention r(13)=0.5861, *p*=0.0353), (VEH %social exploration r(12)= -0.7056, *p*=0.0104) but these associations were not significantly regulated by NIF **(Fig 6C)**. In males, there was a significant negative correlation between %social exploration in Social vs. Object choice and Open Field behavior (r(16)= -0.5209, *p*=0.0385); inhibiting NAc pruning during adolescence had a significant effect on this correlation **(Fig 6D)** (z=-2.49, *p*=0.0128).

**Fig 6.**
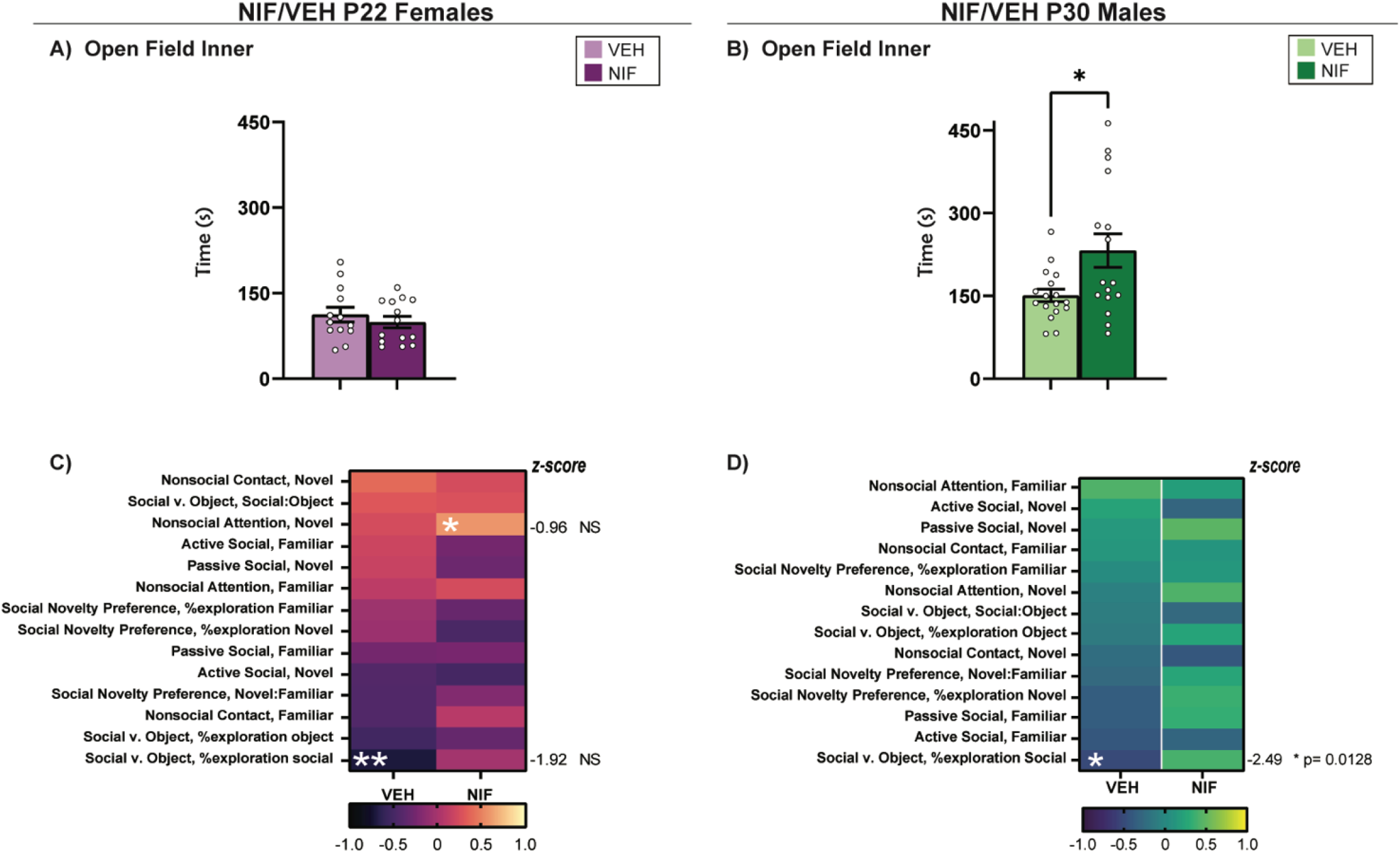
Open Field. (A) After inhibiting NAc pruning during adolescence, females show no statistically significant change in time spent in the inner quadrant of the open field apparatus. (B) In males, there is a statistically significant increase in the amount of time spent in the inner quadrant of the open field apparatus. Histograms represent average ± standard error of the mean. Student’s two-tailed t-test. If ^*^ then p< 0.05, n=13-17. (C) Correlations between open field behavior and other social metrics in females. In NIF-treated females, there is a significant correlation between nonsocial attention in the Novel Interaction test and open field behavior (uncorrected p=0.0353), and in VEH-treated females open field behavior significantly correlates with %social exploration in the Social vs. Object choice test (uncorrected p=0.0104). (D) Associations between open field behavior and other social metrics in males. In VEH-treated males open field behavior correlates to %social exploration in the Social vs. Object choice test (uncorrected p=0.0385). A Fisher’s r-to-z tests indicates this relationship is significantly regulated by NIF treatment and is shifted to a positive (albeit nonsignificant) correlation after NIF treatment (p=0.0128).

## DISCUSSION

These data indicate that a temporally specific manipulation of microglial signaling in the developing NAc will have persistent effects on social behavior. Our hypothesis was that inhibiting NAc pruning during sex-specific adolescent periods would increase sociability in sex-specific ways. To our surprise, we found that this was the case, but sociability was primarily increased toward familiar, and not novel, conspecifics. In males, inhibiting NAc pruning increased familiar exploration in the Social Novelty Preference test and increased nonsocial contact at the expense of nonsocial attention in the Familiar interaction test. In females, inhibiting NAc pruning did not change in Social Novelty Preference behavior, and increased active social interaction at the expense of nonsocial contact in the Familiar Interaction test. This leads us to infer that *naturally occurring* pruning serves to reduce social behaviors primarily directed toward a familiar conspecific in both sexes, but in sex-specific ways.

### 1. Adolescent social development shapes adult behavior

Adolescence is a period characterized by increased risk-taking behavior, heightened novelty seeking, and significant social maturation (Blankenstein et al., 2020; Bukowski & Adams, 2005; Spear, 2000; Sturman & Moghaddam, 2012) These behaviors likely facilitate “leaving the nest,” i.e. the maturation of the dependent child to independent adult. We anticipated broad effects on sociality in both sexes, indiscriminate of the social partner. What we found, however, is that most social changes induced by inhibiting NAc pruning during adolescence were present when a familiar conspecific was incorporated into the social test, and very little change in response to a novel social stimulus. This may suggest that adolescent NAc pruning is less important for the natural increase in peer-centered social behaviors that emerge during adolescence, and more important for reducing familiarity-associated social behaviors. If true, this would also suggest different neural mechanisms guiding familiarity-vs. novelty-directed social development, which to our knowledge is a new concept. Our data suggest it may be important to incorporate familiar stimuli into social tests for more comprehensive assessments of sociability. Another possibility considers the implications of increased rewarding social play during adolescence (Douglas et al., 2004; van den Berg et al., 1999), which we previously published was an acute consequence of inhibiting NAc pruning in both sexes. Social enrichment over adolescence is crucial for “normal” development (Burleson et al., 2016; van den Berg et al., 1999), and adolescent social isolation stress confers risk towards maladaptive behaviors, including anxiety-like (Huang et al., 2017; Lukkes et al., 2009; Weintraub et al., 2010), depression-like (Bukowski & Adams, 2005), and addiction-like behaviors (Deutschmann et al., 2022; Surakka et al., 2021). Interestingly, rats deprived of social play during adolescence have heightened anxiety-like behavior (Parvopassu et al., 2021). Our data complement this finding, as males that had inhibited NAc pruning during adolescence have a lowered anxiety-like phenotype **(Fig. 6B)**. Whether increased social play would strengthen the social relationship between the familiar cage mates, and the consequences of such a bond, remain to be tested. Pair bonding is most often studied in the monogamous prairie vole, between mates (Goodwin et al., 2019; Insel et al., 1998; Lieberwirth & Wang, 2016). Sibling/cage mate bonds have emerged as an area of interest only recently (Lee & Beery, 2021, 2022), but are less commonly studied, irrespective of animal model (Cirulli et al., 1996; Smith et al., 2018). In females, pup-rearing can be used to study social behavior and affiliative bonds (Ahern & Young, 2009). Across species, co-rearing by 2+ females improves reproductive success and survival of the offspring (Bredy et al., 2007; Martinez et al., 2015; Rox et al., 2022). Furthermore, maternal behaviors in rodents, while innate (Okabe et al., 2017; Rincón-Cortés & Grace, 2020), can be socially taught to a nulliparous female (Carcea et al., 2021). Increased active social attention, therefore, in female cage mates **(Fig 5A)** could be evolutionarily advantageous in communal pup-rearing environments. For a number of reasons, increased active social attention is not always positive, especially in males [(Lukas & de Jong, 2017; Van Loo et al., 2003). NIF-treated males do not increase active social attention with a novel or familiar conspecific (**Fig. 4E, Fig. 5E**). What relevance increased social contact has in the NIF-treated males is also an open question (**Fig 5H**) (Bales et al., 2018; Saarinen et al., 2021; Wu et al., 2021).

### 2. Microglial pruning may be a linchpin in healthy and abnormal social development

Neurodevelopmental plasticity is tightly regulated, temporally bound, and can be sex-specific based on the brain region examined (Hanamsagar et al., 2017; Schwarz et al., 2012; Weinhard, Neniskyte, et al., 2018). Microglia, the innate immune cells of the brain, mediate the synaptic pruning process by phagocytosing synaptic proteins (Cangalaya et al., 2020; Weinhard, di Bartolomei, et al., 2018), among larger neural structures (Kurematsu et al., 2022). The complement system is a critical component of peripheral immunity and has more recently been accepted as a driver of development and plasticity (Mastellos, 2014; Presumey et al., 2017), specifically in activity-dependent pruning (Gyorffy et al., 2018; Lieberman et al., 2019; Schafer et al., 2012; Thion & Garel, 2018). Data are now accumulating that microglial pruning plays a role in social development via the regulation of neural development in several different brain regions (Kato et al., 2012; Kopec et al., 2018). Sex differences in microglial pruning can be induced by the prenatal androgen surge (Lenz et al., 2013) and serve to differentiate mature social behaviors based on gonadal sex in the medial preoptic area and amygdala (VanRyzin et al., 2019; VanRyzin et al., 2020). During adolescence, the synaptic landscape of the NAc undergoes a period of neuroimmune plasticity (Matthews et al., 2013; Schramm et al., 2002) coinciding with a peak and decline of social play (Manduca et al., 2016). Interfering with C3 receptor function on microglia appears to increase the length of time of the peak of social play (Kopec et al., 2018). Our data suggest while the short-term consequence of NAc pruning at sex-specific times shows a convergent behavioral phenotype (increased play), the long-term programming of the social brain becomes sex-specifically refined. Moreover, there is evidence that the neurochemical systems being regulated by pruning in the NAc are sex-specific: dopamine signaling (via dopamine D1 receptor pruning) in males and an unknown, non-D1r system in females. The NAc is a heavily innervated region (Bayassi-Jakowicka et al., 2021), and thus the downstream consequences of differential synaptic modulation between the sexes are likely to have wide-reaching effects (Louilot et al., 1986; Stark et al., 2023). The biological and behavioral importance of pruning different neurochemical systems in each sex to regulate similar, but not entirely overlapping social behaviors in adulthood, is unclear at this time. For example, microglia-mediated pre-synaptic excitatory pruning in the prefrontal cortex during adolescence, after the adolescent NAc pruning timepoint, also impacts later-life social behaviors (Andrews et al., 2021; Cressman et al., 2010). There may also be a role for microglia-mediated synaptic pruning in the hippocampus during the juvenile period for social development, but the evidence was collected with a global transgenic mouse with microglial deficiencies, and thus it is difficult to make conclusive statements (Zhan et al., 2014) Finally, while these reports serve as evidence that microglial pruning is important for natural social development, there are also reports suggesting that changes to microglial pruning in early life are involved in the pathogenesis of many neurodevelopmental disorders that have social symptoms, including, autism-spectrum disorder (Arcuri et al., 2017; Kim et al., 2017; Ma et al., 2020; Zhan et al., 2014) and schizophrenia (Boksa, 2012; Howes & McCutcheon, 2017; Jones et al., 2020; Presumey et al., 2017). These data would suggest that understanding the healthy and abnormal regulation of microglia-mediated synaptic pruning, and region-specific effects on social behaviors, may be important to understand complex neurodevelopmental disorders.

### 3. Future directions and limitations

A limitation of this study is the inability to include direct comparisons of sex, as the timing of NIF manipulations was sex specific. Future studies may focus on better understanding sex differences in the consequences of NAc pruning in a more direct way, for both social and nonsocial behaviors. Moreover, our previous work would suggest that the social consequences of inhibiting pruning in the NAc would be D1r-dependent for males, but not females. Whether altered D1r signaling in the NAc can account for all the male changes will be important to determine in future studies. Finally, future studies may include investigation of the molecular landscape of the adult NAc after pruning, to determine long-term effects of pruning on local and global signaling in the NAc.

In conclusion, the data presented in this report indicate that interfering with a microglia-mediated developmental process within a single brain region during adolescence is sufficient to alter social developmental trajectories in sex-specific ways.

## Acknowledgements

This work was supported by the National Institutes of Health R01DA052889 and R03AG07011 to AMK and Albany Medical College Start-up funds to AMK.

## Author Declarations

JMK and AMK designed the experiments. JMK performed the experiments. All authors analyzed the experiments. JMK and AMK wrote the manuscript. All authors edited the manuscript. The authors declare no conflicts of interest.

## FIGURE LEGENDS

**Supp. Fig. 1.**
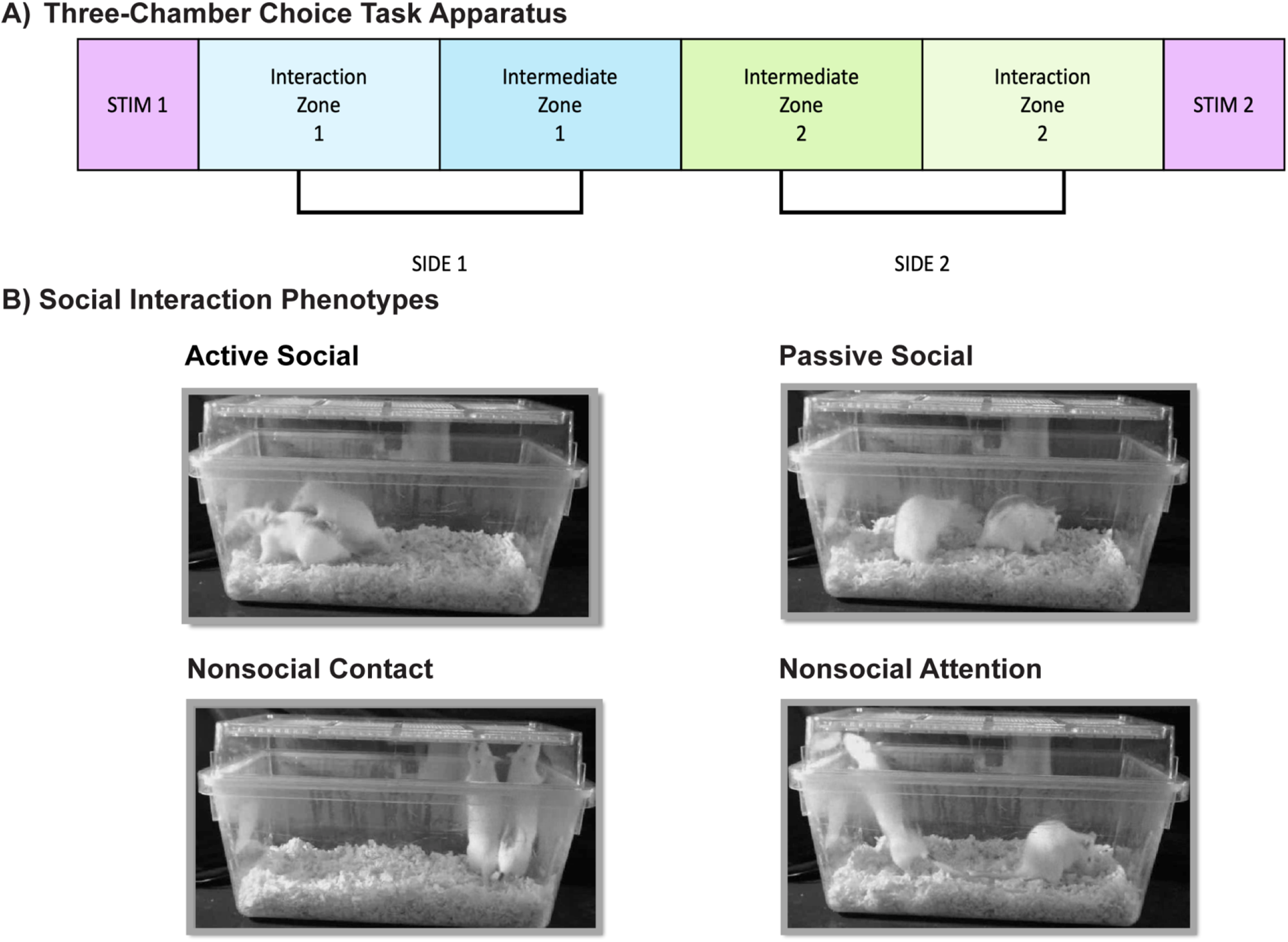
(A) Schematic of choice tasks apparatus. %exploration= time in interaction zone/time in target side. (B) Representative images of interaction metrics

